# Comparative Genomics of COBRA-like Genes in *Theobroma* and *Herrania* Reveals Structural Conservation and Lineage-Specific Variation

**DOI:** 10.64898/2026.09.14.751154

**Authors:** Rafaely Pantoja Oliveira, João Victor dos Anjos Almeida, Vitor Trinca, Anelise Stella Ballaben, Mauro de Medeiros Oliveira, Alessandro de Mello Varani

## Abstract

COBRA-like (*COBL*) genes encode plant-specific proteins associated with cell-wall organization, cellulose deposition, and developmental processes. Despite their functional relevance in model plants and crops, this gene family remains poorly characterized in Amazonian Malvaceae species. Here, we performed a comparative genomic analysis of *COBL* genes in two *Theobroma grandiflorum* clones, two *T. cacao* cultivars, and *Herrania umbratica*. Using homology searches, conserved domain validation, phylogenetic reconstruction, gene structure analysis, chromosomal mapping, transcript abundance profiling, molecular modeling, and selection tests, we identified 62 *COBL* genes across the analyzed genomes. *COBL* copy number varied among genomes, with *T. cacao* Matina showing the largest repertoire. All retained proteins contained the conserved COBRA domain and grouped into the two major COBL subgroups previously described in angiosperms, with subgroup-specific exon–intron organization largely conserved across species. Genomic distribution and duplication classification indicated contributions from tandem, segmental, dispersed, and proximal duplications, suggesting that multiple genomic processes shaped COBL family organization in *Theobroma* and *Herrania*. Phylogenetic analyses further recovered a putative lineage-specific COBL clade within the analyzed Malvaceae, indicating lineage-specific variation in this gene family. Structural modeling indicated overall conservation between representative proteins from this clade and *COBL6*- associated proteins, although localized amino acid differences were observed within the COBRA domain. Selection analyses supported predominant purifying selection across the family, with limited evidence of episodic diversifying selection in specific lineages and no significant branch-level signal within the putative lineage-specific clade after correction. Overall, these results provide a comparative framework for *COBL* gene evolution in *Theobroma* and *Herrania* and identify candidate genes for future functional studies on cell-wall-related traits in economically important Malvaceae crops.

**Highlights:**

- Sixty-two COBRA-like genes were identified across *Theobroma* and *Herrania* genomes.
- *COBL* genes retained conserved subgroup architecture despite genome-specific copy number variation.
- Multiple duplication modes shaped *COBL* diversification in the analyzed Malvaceae.
- A putative Malvaceae-associated *COBL* lineage was recovered in *Theobroma* and *Herrania*.
- *COBL* genes evolved predominantly under purifying selection.

## 1. Introduction

*Theobroma grandiflorum* (Willd. ex Spreng.) K.Schum., commonly known as cupuassu, and *T. cacao* L. are Neotropical tree species of the family Malvaceae. Among the 23 cataloged species of *Theobroma* L., cupuassu and cacao are notable for their agricultural, commercial, and cultural importance [1–3]. The closely related species *Herrania umbratica* produces berry-like fruits with sweet pulp, consumed by humans and wildlife [1,4], and provides a useful comparative lineage for genomic analyses involving *Theobroma*. Although recent molecular studies have suggested a close relationship between *Theobroma* and *Herrania*, including proposals to synonymize *Herrania* within *Theobroma*, this taxonomic interpretation remains under discussion, and both genera are still commonly treated as distinct lineages [1,2,5].

Cacao is one of the most economically important species of *Theobroma*, largely due to the global demand for chocolate and other cacao-derived products. In the Amazonian context, cupuassu has also gained economic relevance through its use in juices, jams, sweets, ice creams, butters, cosmetics, and the chocolate-like product known as “cupulate” [6–10]. In Pará, the main producing state in Brazil, cupuassu cultivation occupies approximately 8.9 thousand hectares, with an annual production of about 29 thousand tons [11,12]. Its regional importance, particularly for small-scale farmers and agroforestry systems, supports the development of genomic studies that may contribute to future breeding and crop improvement strategies [13].

Cell-wall formation and remodeling are central processes in plant growth, development, tissue organization, and interactions with the environment. In fruit crops, cell-wall-related mechanisms may influence traits such as texture, firmness, postharvest behavior, and pathogen responses. Therefore, identifying gene families involved in cell-wall biology can provide useful candidates for future functional studies in economically relevant crops. In cupuassu and cacao, the availability of genomic resources now enables comparative analyses of gene families potentially associated with these traits [14–17].

Among cell-wall-associated gene families, the COBRA-like (COBL) family is of particular interest. This multigene family, homologous to the COB gene in *Arabidopsis thaliana*, encodes plant-specific glycosylphosphatidylinositol (GPI)-anchored proteins that typically contain an N-terminal secretion signal and a C-terminal region with an ω-site for GPI modification [18,19]. COBL proteins are characterized by the conserved COBRA domain (PFAM: PF04833), which has been associated with cell-wall organization and cellulose deposition [20]. COBL proteins have been implicated in diverse physiological and developmental processes, including stem mechanical strength, pollen tube elongation, pathogen responses, and root hair formation [9,18,19,21,22].

The *COBL* gene family is evolutionarily conserved across monocots and eudicots [20,23], with homologs described in species such as tomato, maize, cotton, black cottonwood, rice, and *A. thaliana* [20,24–28]. Previous studies have classified COBL proteins into two major subgroups distinguished by structural features, including differences in protein organization and exon–intron architecture [20]. However, despite their biological relevance in model and crop species, *COBL* genes remain poorly characterized in *Theobroma* and closely related genera.

Here, we performed a comparative genomic analysis of *COBL* genes in two *T. grandiflorum* clones, two *T. cacao* cultivars, and *H. umbratica*. Using homology searches, conserved domain validation, phylogenetic reconstruction, gene structure analysis, chromosomal mapping, transcript abundance profiling, molecular modeling, and selection tests, we investigated the evolution and structural conservation of this gene family in *Theobroma* and *Herrania*. Rather than directly assigning biological functions, this study provides a comparative framework for *COBL* gene evolution and identifies candidate genes for future functional studies on cell-wall-related traits in economically important Malvaceae crops.

## 2. Materials and methods

### 2.1. Genomes selected for the study

This study used genomic data from two *Theobroma grandiflorum* clones: C1074, susceptible to witches’ broom disease, and C174, also known as Coari, resistant to witches’ broom disease [29,30]. Both samples belong to the Embrapa Amazônia Oriental genetic collection, located in Belém, Pará, Brazil [16,31]. In addition, reference genomes of *T. cacao* cultivars Criollo and Matina, assembled at chromosomal scale, and the draft genome of *Herrania umbratica* cultivar Fairchild were included for comparative analyses [32,33]. Genomic sequences and annotation files were retrieved from publicly available records in GenBank under the accession numbers PRJNA383741, PRJEB14326, and PRJNA239715, respectively. These datasets are also integrated into the Plant Genomics data platform (https://plantgenomics.ncc.unesp.br), which provides a genome browser and user-friendly tools for browsing genomic annotations and gene models. Therefore, all gene identifiers reported in this study follow the gene ID nomenclature adopted in the Plant Genomics platform.

### 2.2. Identification and Chromosomal Localization of COBRA-like Gene Family Members

Protein sequences corresponding to COBRA-like gene family members in *A. thaliana* were obtained from the UniProt database and used as reference queries: *COB* [Q94KT8], *COBL1* [Q9SRT7], *COBL2* [Q8L8Q7], *COBL4* [Q9LFW3], *COBL5* [Q9FME5], *COBL6* [O04500], *COBL7* [Q8GZ17], *COBL8* [Q9LIB6], *COBL9* [Q9FJ13], *COBL10* [Q9LJU0], and *COBL11* [Q9T045] (Table 1). These sequences were used to identify similar predicted proteins in the annotation files of *T. grandiflorum* C174 and C1074, *T. cacao* Criollo and Matina, and *H. umbratica*.

**Table 1.**
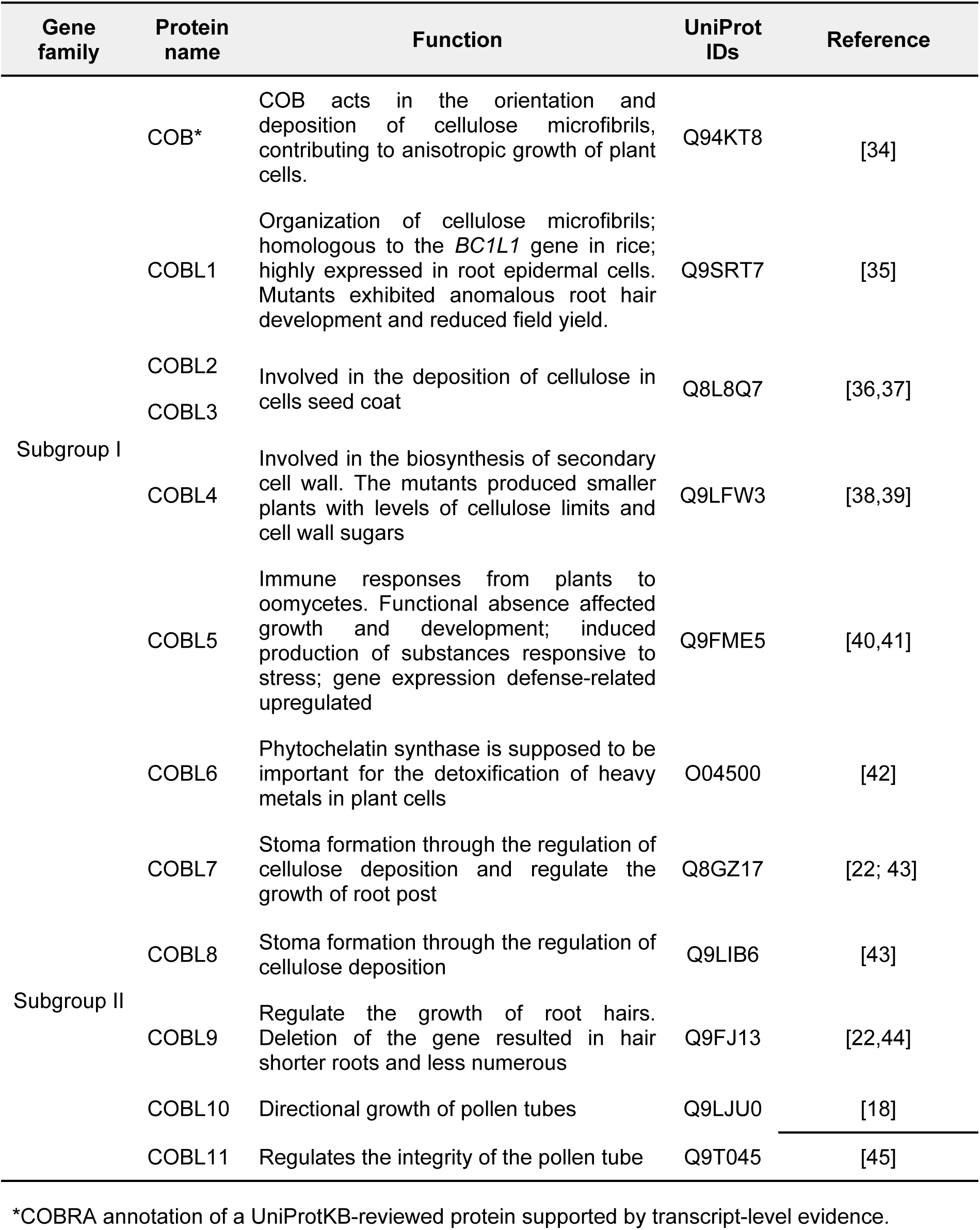
COBRA-Like gene family members in the *Arabidopsis thaliana* genome play essential roles in cellulose biosynthesis, crystallization, and integration into the cell wall.

Candidate COBL proteins were identified through sequence similarity searches performed on the Plant Genomics platform. Initial candidates were retained using an E-value threshold of 1e-5 and minimum sequence similarity and alignment coverage values of 60%. The presence of the conserved COBRA domain (PF04833) was subsequently evaluated using the HMMER web server v2.41.2 [46] in combination with the Pfam database [47]. To reduce false-positive assignments, only proteins containing the COBRA domain were retained for downstream comparative analyses.

*COBL* genes were mapped onto the chromosomes of *T. grandiflorum* and *T. cacao*, and onto scaffolds of *H. umbratica*, using gene coordinates obtained from the corresponding genome annotation files. Chromosomal and scaffold distribution maps were generated using MG2C v2.1 [48].

### 2.3. Transcript Abundance Analysis of COBRA-like Genes

For *T. grandiflorum*, transcript abundance analyses were performed using young leaf RNA-seq data previously generated by Alves et al. [29] and de Abreu et al. [30]. In those studies, total RNA was extracted from fresh leaves using a modified CTAB protocol [49], and Iso-Seq data were also generated to support genome annotation. In the present study, these previously generated transcriptomic resources were used to assess *COBL* transcript abundance and annotation support in *T. grandiflorum*.

For *H. umbratica* and *T. cacao*, publicly available RNA-seq datasets from different plant tissues were retrieved from GenBank. The *H. umbratica* datasets included lateral buds, closed flower buds, apical stems, young leaves, and open flowers. The *T. cacao* datasets included leaves, pod husks, stems, roots, flowers, and seeds. Transcript abundance was estimated as TPM, and the GenBank SRA accession numbers for all RNA-seq datasets used in these analyses are provided in Supplementary Material 2.

Because RNA-seq datasets differed among species in tissue composition, experimental origin, and sampling design, TPM values were interpreted as exploratory evidence of transcript detection rather than as direct quantitative comparisons among species.

### 2.4. Gene Structure, Conserved Domain, and Duplication Analyses

Structural information on exons, introns, untranslated regions (UTRs), gene coordinates, and gene duplication classes for the analyzed genomes was retrieved from the genomic annotations and comparative genomic analyses previously generated by Alves et al. [29] and de Abreu et al. [30], and made available through Plant Genomics DB. These resources include gene models and duplication classifications inferred using MCScanX, commit b1ca533 [50], which classifies genes according to genomic topology into whole-genome or segmental duplication, tandem duplication, proximal duplication, and dispersed duplication categories.

For the present study, *COBL* genes identified in the analyzed genomes were associated with their corresponding structural annotations and MCScanX-based duplication classes available in Plant Genomics DB. Exon–intron organization and conserved protein domains were analyzed and visualized using TBtools v2.019 [51]. These data were used to describe the structural organization and duplication patterns of COBL family members across the analyzed *Theobroma* and *Herrania* genomes.

### 2.5. Phylogenetic Analysis of the COBRA-like Gene Family

Phylogenetic relationships among COBL proteins from *T. grandiflorum*, *T. cacao*, *H. umbratica*, and *A. thaliana* were inferred using the Maximum Likelihood (ML) method [52]. Complete COBL protein sequences from *A. thaliana* were retrieved from the UniProt database and included to support comparative classification of COBL subgroups.

Multiple sequence alignments of COBL protein sequences were generated using MAFFT v7 [53]. Phylogenetic trees were constructed using IQ-TREE v2 [54]. The best-fit substitution model was selected using the Bayesian Information Criterion (BIC) implemented in ModelFinder [55], and JTT+G4 was selected as the best model for the main COBL dataset. Branch support was assessed using 1,000 ultrafast bootstrap replicates. Phylogenetic trees were visualized using iTOL v6 [56].

The phylogenetic analysis was further expanded to include COBL protein sequences from species representing the orders Asterales, Brassicales, Cucurbitales, Fabales, Fagales, Lamiales, Malpighiales, Malvales, Poales, Rosales, and Solanales (Supplementary Material 3). This expanded dataset was used to evaluate whether COBL sequences from *T. cacao*, *T. grandiflorum*, and *H. umbratica* clustered with COBL proteins from other angiosperm lineages. For this dataset, alignments were also generated using MAFFT, and phylogenetic inference was performed using IQ-TREE. The best-fit model selected according to BIC was JTTDCMut+R10.

Genomic data for the expanded analysis were obtained from NCBI RefSeq and GenBank. The presence of the COBRA domain in the proteome of each species was confirmed using InterProScan v5.67-99.0 [57]. To standardize comparisons across annotation datasets, redundant isoforms were removed by retaining the longest protein isoform for each gene.

### 2.6. Selection Analysis

Episodic selection was evaluated using HyPhy v2.5.94 (https://www.hyphy.org/). Codon-aware alignments were generated with PAL2NAL [58] using the MAFFT protein alignment of Malvaceae COBL sequences and their corresponding coding sequences. Evidence of episodic diversifying selection at the branch level was tested using the adaptive Branch-Site Random Effects Likelihood model (aBSREL; [59]), considering branches with ω > 1 and corrected p-values ≤ 0.05 as significant. Multiple-test correction was performed using the procedure implemented in HyPhy.

Site-level episodic selection was evaluated using the Mixed Effects Model of Evolution (MEME; [60]), with significance assessed using p ≤ 0.05. Posterior probability ≥ 0.80 was used as an additional criterion to support site-level signals when applicable. Selection analysis outputs were visualized using HyPhy Vision (https://vision.hyphy.org/), and final images were edited in Inkscape.

### 2.7. Molecular Modeling and Structural Analysis of COBRA-like Proteins

Three-dimensional structures of selected COBL proteins encoded by genes from the analyzed species were predicted using the AlphaFold v3 platform [61]. The stereochemical quality of the predicted structures was evaluated using the SAVES 6.0 platform (https://saves.mbi.ucla.edu/), including Ramachandran plot analysis to assess the distribution of residues across conformational regions [62]. For structural comparison, representative COBL proteins from the putative lineage-specific clade and from the COBL6-associated group were selected. COBL6-associated proteins were included because this group represented the closest phylogenetic comparison to the putative lineage-specific COBL proteins and shared similar exon organization. Protein structure files initially generated in CIF format were converted to PDB format using PyMOL v2.3.1 (https://www.pymol.org/).

Structural superposition was conducted using FATCAT v2.0, which performs flexible structural alignments and allows comparison of protein structures without imposing rigid-body constraints [63]. This approach was used to compare overall structural similarity and localized differences along the protein chain. The structural superposition analysis was specifically performed using the proteins TgrandC1074G00000008413.1 and TgrandC1074G00000008646.1. TgrandC1074G00000008646.1 was selected as a representative COBL protein from the putative lineage-specific clade, whereas TgrandC1074G00000008413.1 was selected as a representative COBL6-associated protein from the same genome. These proteins were selected to provide a controlled comparison between proteins from the same genomic background. Other proteins from these groups showed broadly similar predicted structural patterns during preliminary inspection; therefore, these two proteins were used as representative models for structural comparison.

## 3. Results

### 3.1. Identification, Genomic Distribution, and Expression Patterns of COBRA-like Genes in *T. grandiflorum*, *T. cacao*, and *H. umbratica*

A total of 62 genes encoding COBL proteins were identified across the analyzed *T. grandiflorum*, *T. cacao*, and *H. umbratica* genomes (Supplementary Material 4). Among the analyzed genomes, *T. cacao* Matina showed the highest number of *COBL* genes (n = 16), followed by *T. cacao* Criollo and *T. grandiflorum* C1074 (n = 12 genes each), *H. umbratica* (n = 12 genes), and *T. grandiflorum* C174 (n = 10 genes). In each analyzed species or clone, one *COBL* gene was recovered outside the main groups defined by the *A. thaliana COBL* reference sequences, indicating the presence of a putative lineage-specific COBL group in the analyzed Malvaceae genomes.

The genomic distribution of *COBL* genes varied among the analyzed genomes (Fig. 1). In *Theobroma*, *COBL* genes were unevenly distributed across chromosomes. No *COBL* genes were detected on chromosomes 5, 8, 9, and 10 in *T. cacao* Criollo, *T. cacao* Matina, or *T. grandiflorum* C1074. In *T. grandiflorum* C174, no *COBL* genes were detected on chromosomes 5, 7, 8, 9, and 10. In *H. umbratica*, whose genome is currently available as a draft assembly, *COBL* genes were identified on five scaffolds: 5, 8, 9, 11, and 14.

**Fig. 1.**
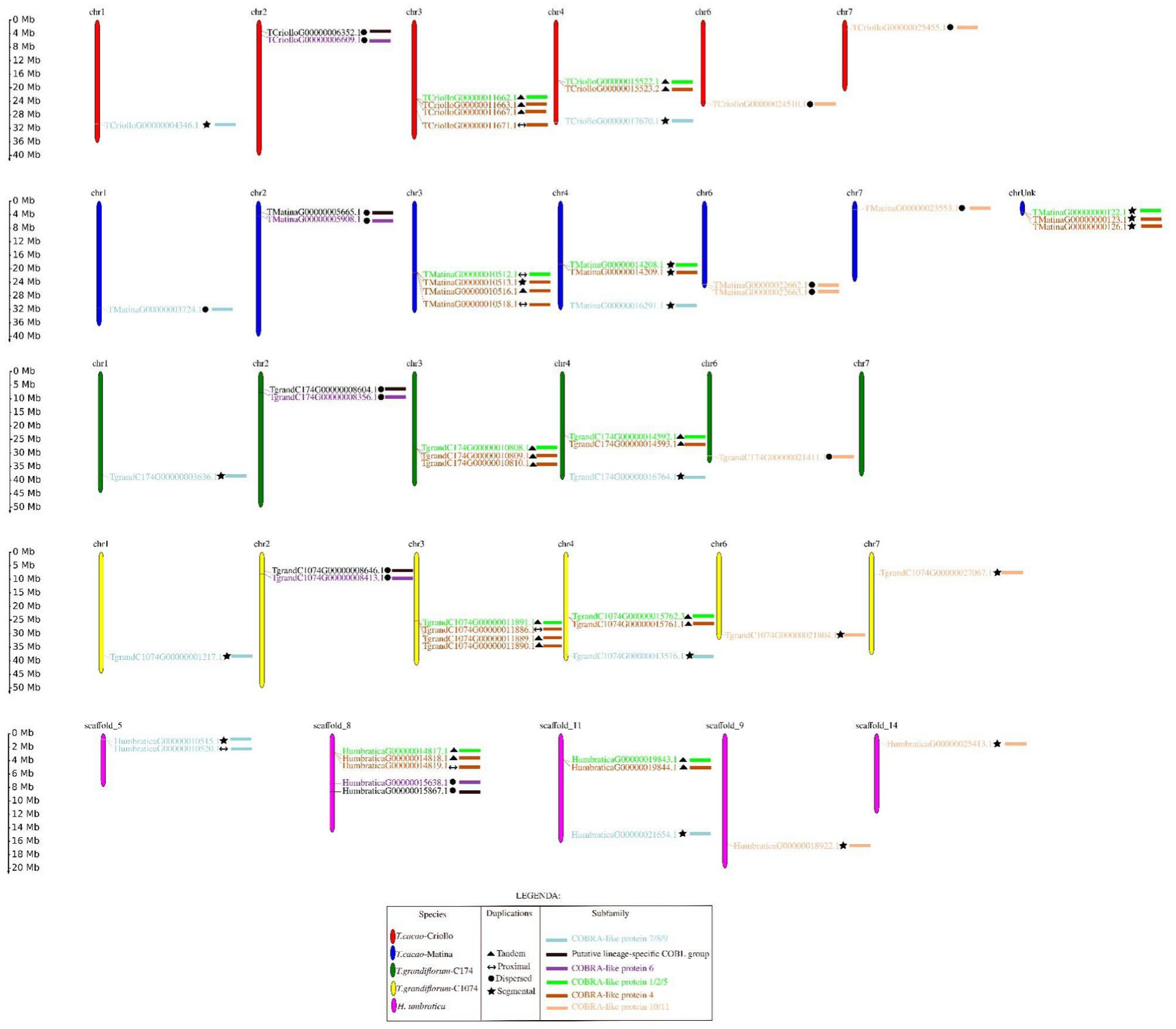
Chromosomal and genetic mapping of the identified *COBL* genes, outlining their specific locations in the studied genomes.

In *Theobroma*, *COBL* genes were mainly concentrated on chromosomes 3 and 4, which together contained 54% of the *COBL* genes identified in this genus. In *H. umbratica*, scaffolds 8 and 11 contained the highest proportion of *COBL* genes, together representing 66.7% of the identified *COBL* repertoire. Duplication classification indicated that tandem duplications were the most frequent category in *T. grandiflorum* C174, *T. grandiflorum* C1074, and *T. cacao* Criollo, whereas segmental duplications were more frequent in *T. cacao* Matina (Table 2). In *H. umbratica*, tandem and segmental duplications occurred at similar frequencies. Proximal and dispersed duplications were less frequent in most analyzed genomes, although dispersed duplications were relatively more represented in *T. cacao* Matina and Criollo (Table 2).

**Table 2.** Number of *COBL* genes derived from different types of gene duplication in *T. grandiflorum* (clones C174 and C1074), *T. cacao* (cultivars Matina and Criollo) and *H. umbratica*. Types of duplication include tandem, proximal, dispersed, and large-scale (segmental).

| Duplication | C174 | C1074 | Matina | Criollo | *H. umbratica* |
| --- | --- | --- | --- | --- | --- |
| Tandem | 5 | 5 | 1 | 5 | 4 |
| Proximal | 0 | 1 | 2 | 1 | 2 |
| Dispersed | 3 | 2 | 6 | 4 | 2 |
| Segmental | 2 | 4 | 7 | 2 | 4 |

Transcript abundance analysis showed that most *COBL* genes had detectable expression in at least one of the analyzed RNA-seq datasets (Supplementary Material 5). However, no transcript abundance was detected for TgrandC174G00000021411.1, TgrandC174G00000010810.1, TgrandC1074G00000011889.1, Tcacao-CriolloG00000011671.1, and TcacaoMatinaG00000010518.1 under the sampled conditions. Because the RNA-seq datasets differed among species in tissue composition and experimental origin, these expression profiles should be interpreted as exploratory evidence of transcript detection rather than as direct quantitative comparisons among species. The absence of detectable expression under the analyzed conditions does not, by itself, indicate loss of function.

### 3.2. Gene structure and conserved domain organization of COBRA-like genes

Gene structure and conserved domain analyses were performed to characterize the organization of *COBL* genes in the analyzed genomes (Fig. 2). Phylogenetic comparison with *A. thaliana COBL* reference sequences supported the classification of the identified genes into previously described COBL groups (Fig. 2a). Exon–intron structure varied among COBL groups, with the number of exons ranging from two to six across the analyzed genes (Fig. 2b).

**Fig. 2.**
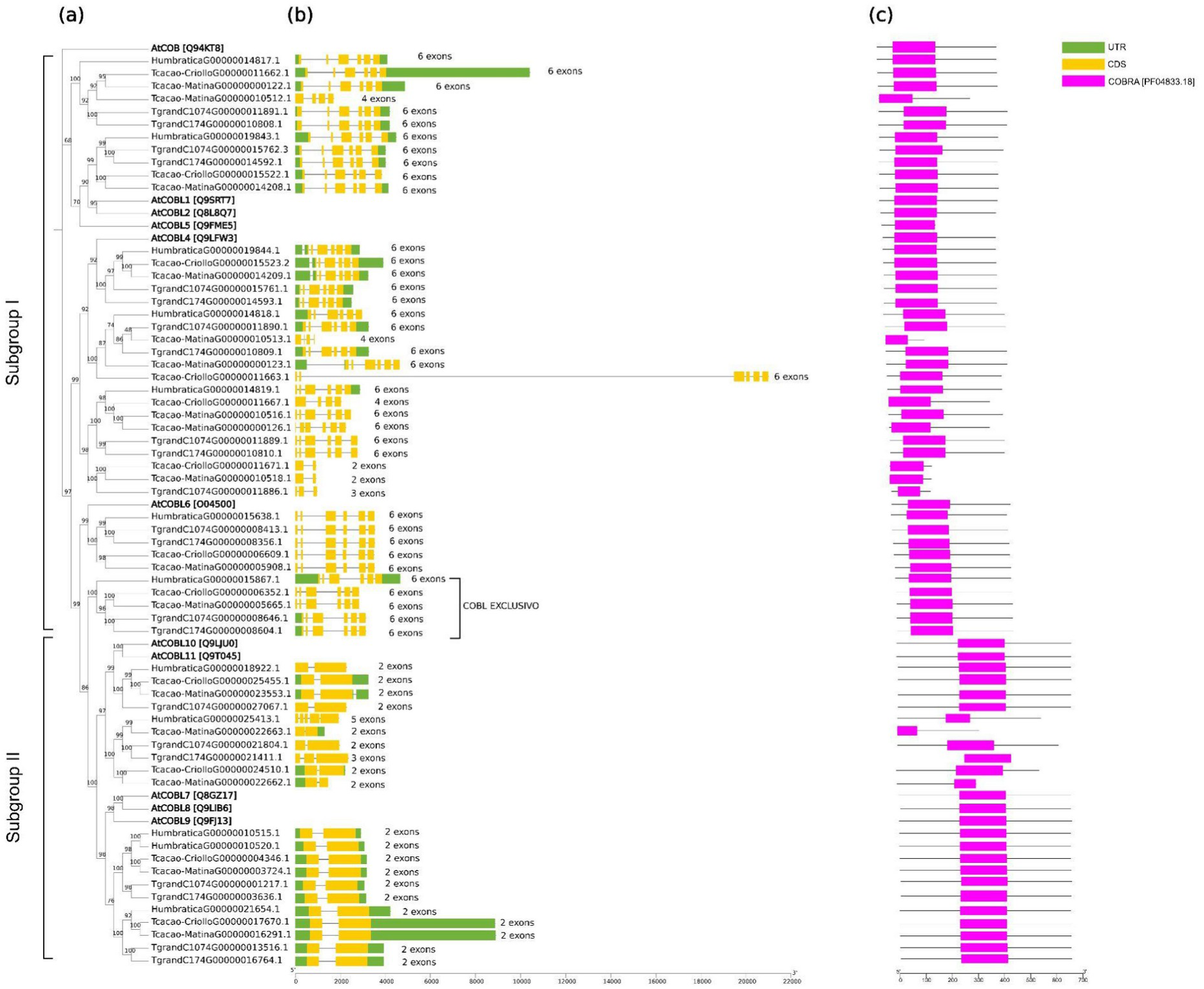
Characterization of the structure and domain of COBL proteins. (a) Phylogenetic tree constructed using Maximum Likelihood method based on 1,000 ultrafast bootstrap trees. The phylogenetic tree was rooted at the midpoint. (b) gene structure, and (c) analysis of the conserved COBRA-like domain.

Genes assigned to the COBL7, COBL8, and COBL9 groups uniformly contained two exons. Most genes assigned to the COBL10 and COBL11 groups also contained two exons, with the exception of TgrandC174G00000022328.1, which contained three exons, and HumbraticaG00000025413.1, which contained five exons. Genes assigned to the putative lineage-specific group, COBL6, and COBL1/2/5 groups generally contained six exons, except for Tcacao-MatinaG00000010512.1, which contained four exons. The COBL4 group, which showed the largest number of representatives in several genomes, exhibited greater variation in exon numbers, ranging from two to six exons, although most *COBL4* genes contained six exons.

Conserved domain analysis confirmed the presence of the COBRA domain (PF04833.18) in all identified COBL proteins from *T. grandiflorum* C174 and C1074, *T. cacao* Criollo and Matina, and *H. umbratica* (Fig. 2c). This domain-based validation supported the assignment of these proteins to the COBRA-like family.

### 3.3. Phylogenetic Analysis of the COBRA-like Gene Family

Phylogenetic analysis of COBL proteins from *T. grandiflorum*, *T. cacao*, *H. umbratica*, and *A. thaliana* recovered the two major COBL subgroups previously described in angiosperms [20] (Fig. 3). Subgroup I included COBL1, COBL2, COBL4, COBL5, and COBL6-related proteins, whereas Subgroup II included COBL7, COBL8, COBL9, COBL10, and COBL11-related proteins. The overall topology was consistent with the structural differences observed among COBL groups, particularly the broader exon number variation in Subgroup I and the more compact exon–intron organization observed in several Subgroup II members.

**Fig. 3.**
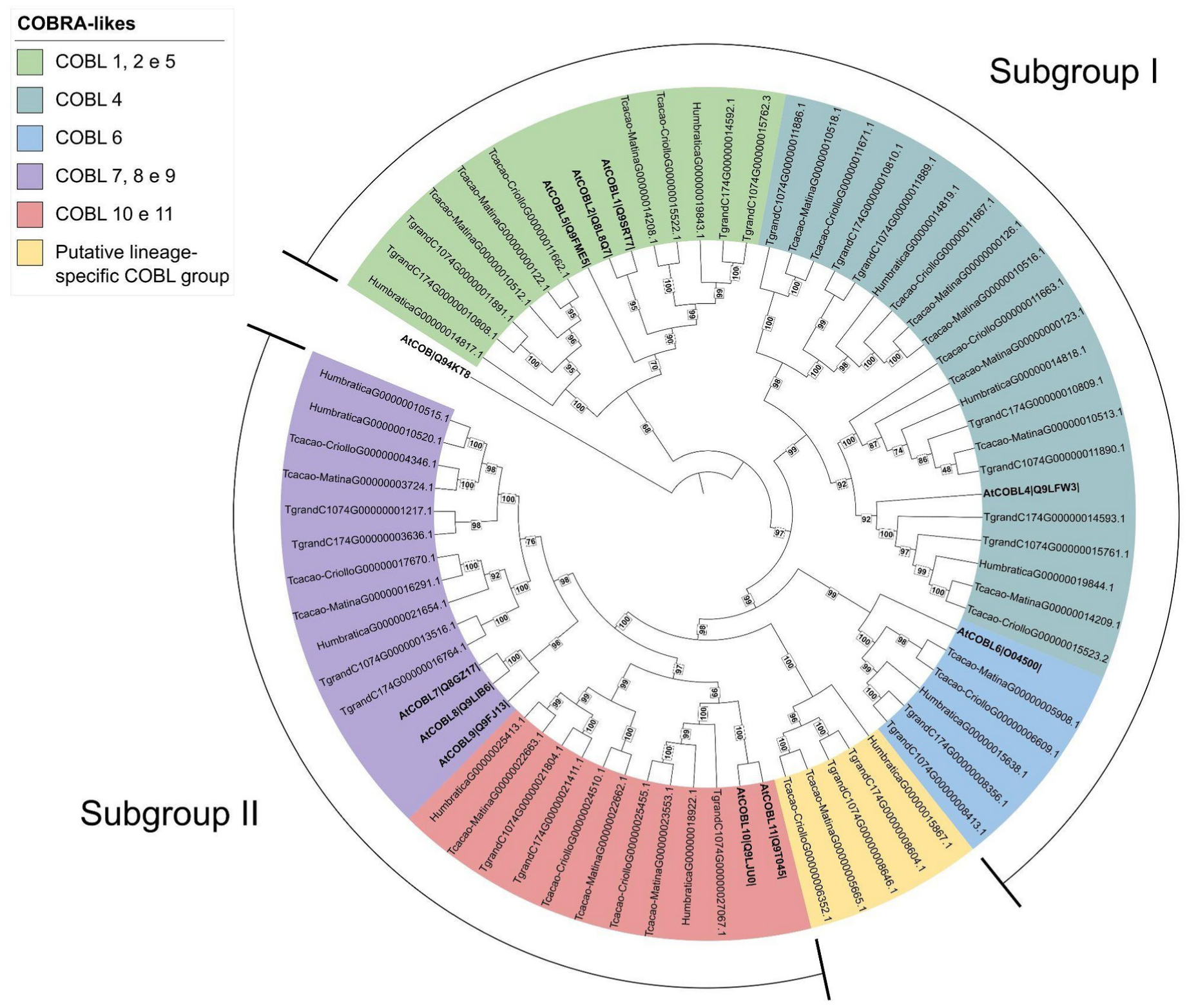
Phylogenetic tree of the COBRA-like family constructed using Maximum Likelihood method.

In addition to the main COBL groups represented by *A. thaliana* reference sequences, the phylogenetic analysis recovered a putative lineage-specific COBL clade containing one representative gene from each analyzed *Theobroma* and *Herrania* genome (Fig. 3). These sequences did not cluster directly with the *A. thaliana* COBL reference groups in the main phylogenetic analysis. To further evaluate the placement of this group, an expanded phylogenetic analysis was performed using COBL proteins from species representing Asterales, Brassicales, Cucurbitales, Fabales, Fagales, Lamiales, Malpighiales, Malvales, Poales, Rosales, and Solanales (Supplementary Material 3; Fig. 4a). In this broader context, the putative lineage-specific COBL sequences from *T. grandiflorum*, *T. cacao*, and *H. umbratica* were recovered within a clade associated with the analyzed Malvaceae/Byttnerioideae sequences and did not cluster directly with COBL sequences from the other sampled angiosperm lineages (Fig. 4b).

**Fig. 4.**
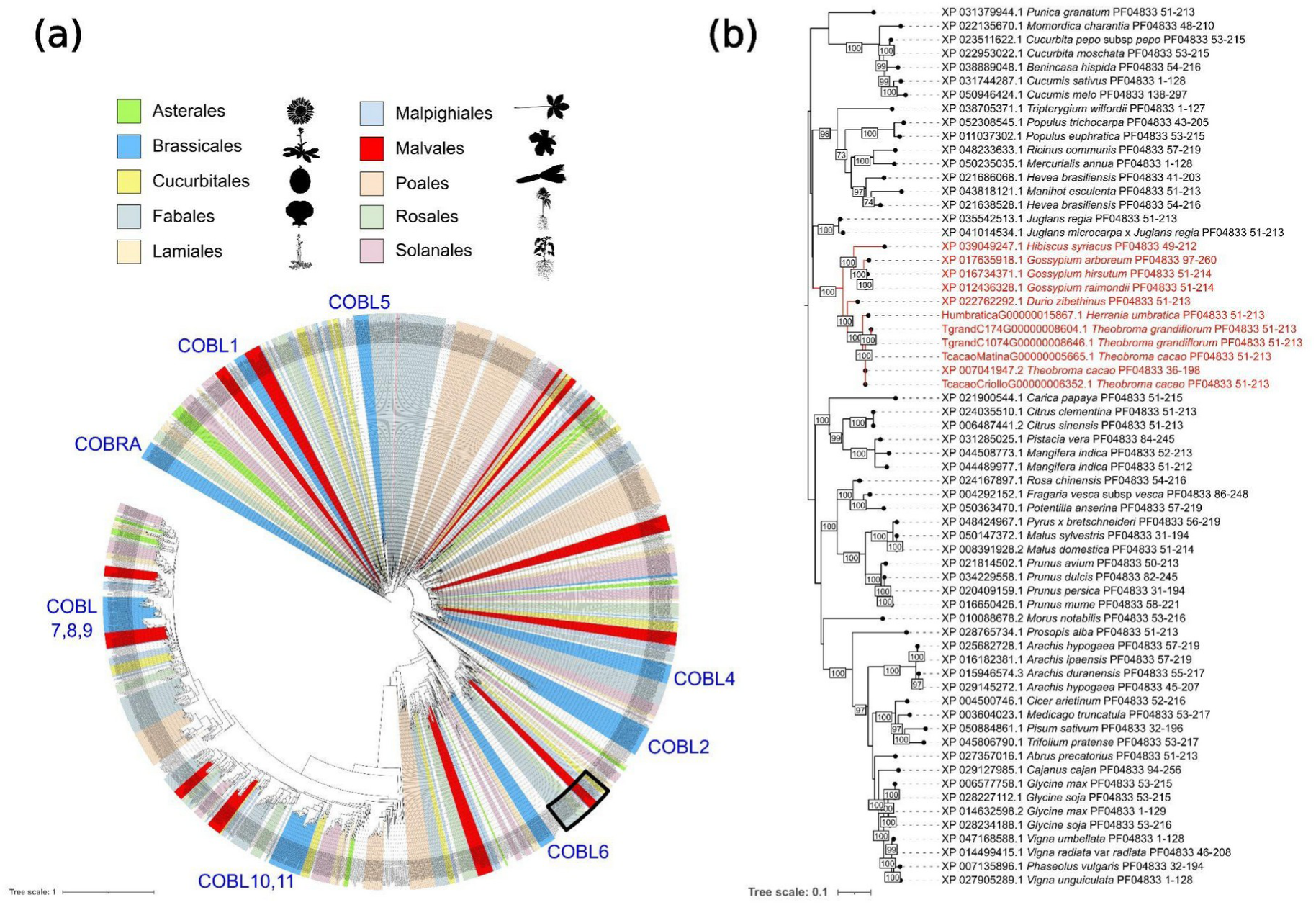
(a) Phylogenetic analysis of COBL proteins from species of the orders Asterales, Brassicales, Cucurbitales, Fabales, Fagales, Lamiales, Malpighiales, Malvales, Poales, Rosales, and Solanales. The phylogenetic tree was rooted using the COBRA protein sequence (Q94KT8) from *Arabidopsis thaliana*. COBL groups are indicated in blue, and the positions of the *A. thaliana* reference sequences among other Brassicales. In (b), a pruned cladogram of the complete tree (black rectangle) is shown, with sequences from the putative lineage-specific COBL group identified in this study highlighted in red on the branches, alongside other COBRA-like domain–containing genes. Silhouettes were obtained from https://www.phylopic.org/.

### 3.4. Structural comparison of representative COBRA-like proteins

Three-dimensional structural models were generated for selected COBL proteins to compare representative proteins from the putative lineage-specific clade and from the COBL6-associated group. The structural comparison focused on TgrandC1074G00000008646.1, representing the putative lineage-specific clade, and TgrandC1074G00000008413.1, representing a COBL6-associated protein from the same genome.

The predicted protein models showed broad structural similarity between these two representative proteins. Predicted proteins corresponding to the products of the COBRA-like genes (Fig. 5a) showed broad structural similarity between the two representative models (Fig. 5b; Supplementary Fig. 1; Supplementary Material 6). Flexible structural alignment using FATCAT also indicated substantial overlap between TgrandC1074G00000008646.1 and TgrandC1074G00000008413.1, with 402 equivalent aligned positions and an RMSD of 1.96 Å (Fig. 5c; Supplementary Material 6). These results indicate overall structural conservation between the representative proteins analyzed.

**Fig. 5.**
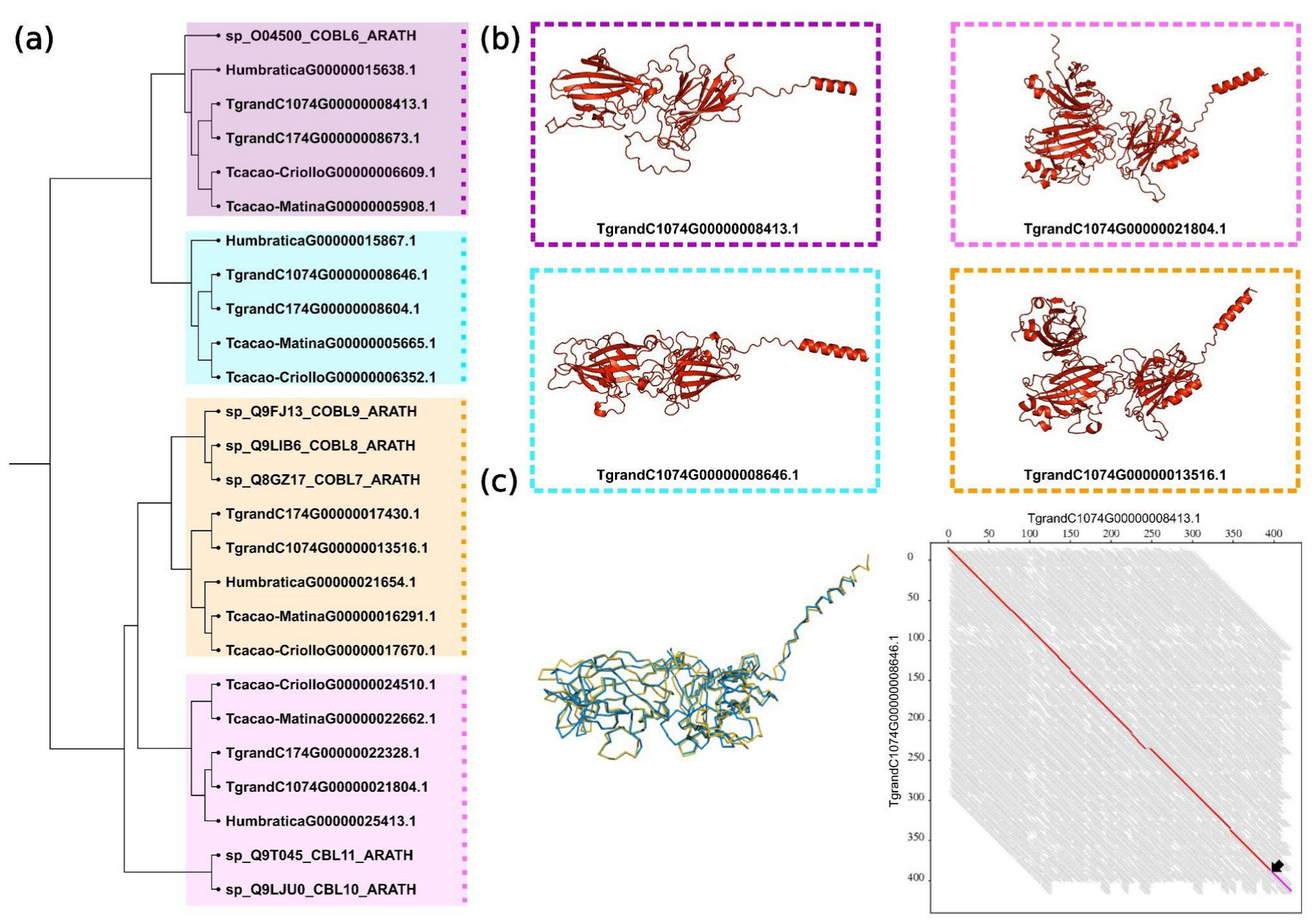
(a) Cladogram showing the relationships among predicted COBL proteins identified in *T. grandiflorum*, *T. cacao*, and *H. umbratica*, clustered together with the COBL6, COBL7, COBL8, COBL9, COBL10, and COBL11 proteins from *A. thaliana*. Colored boxes highlight the major COBL clades identified across the analyzed species. (b) Predicted threedimensional structures of representative COBL proteins from the COBL6-related, COBL7/8/9-related, COBL10/11-related, and the putative lineage-specific COBL clades identified in *T. grandiflorum* C1074. Despite sequence divergence among groups, the predicted models retained a conserved overall fold characterized by β-sheet-rich domains and a C-terminal α-helical region. (c) Structural superposition of the *T. grandiflorum* C1074 proteins TgrandC1074G00000008413.1 and TgrandC1074G00000008646.1 generated using FATCAT, revealing high structural similarity between the proteins (402 aligned positions; RMSD = 1.96 A; P-value = 0). The arrow indicates a flexible twist detected during alignment, corresponding to a local rotation/translation between structural segments

Amino acid sequence comparisons showed localized differences within the COBRA domain between COBL6-associated and putative lineage-specific proteins (Supplementary Fig. 2 and 3). These substitutions identify candidate positions that may be useful for future functional analyses. However, the structural and sequence comparisons alone do not establish the functional impact of these amino acid differences.

### 3.5. Selection analyses of COBRA-like genes

Selection analyses indicated that the *COBL* gene family is predominantly conserved across the analyzed Malvaceae genomes. The overall pattern was consistent with strong purifying selection across most branches. Branch-level analysis using aBSREL identified a limited number of branches with evidence of episodic diversifying selection after correction, including specific lineages such as TgrandC1074G00000021804.1 and Tcacao-MatinaG00000010512.1 (Fig. 6). Most tested branches showed no significant evidence of episodic diversifying selection.

**Fig. 6.**
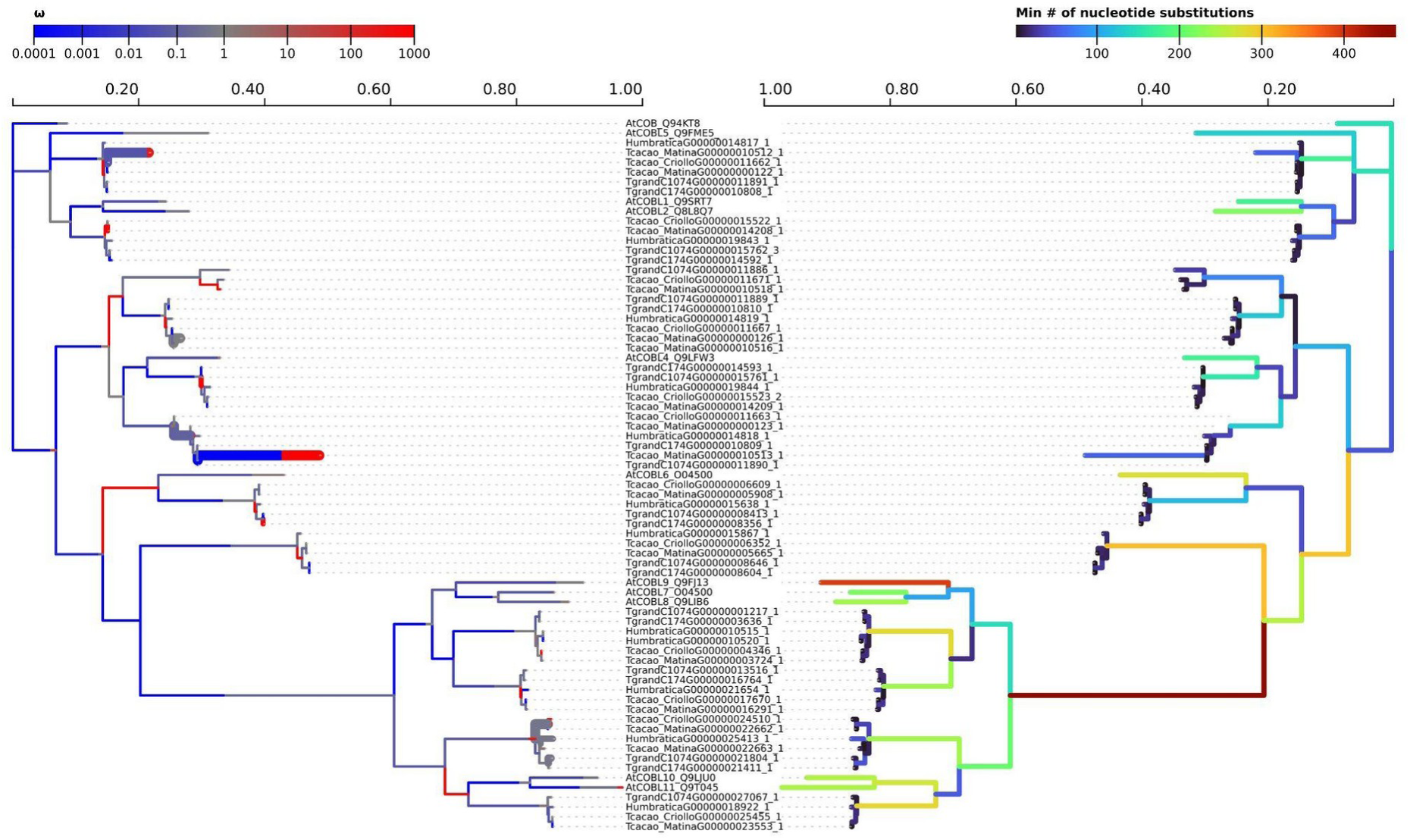
Branch-level episodic diversifying selection in the COBRA-Like gene family inferred using aBSREL. The phylogenetic tree shows COBRA-Like sequences from *T. grandiflorum*, *T. cacao*, *H. umbratica*, and reference COBL proteins from *A. thaliana*. Branch colors represent the estimated branch-site ω values visualized in Hyphy Vision, with branches showing significant evidence of episodic diversifying selection indicated according to the aBSREL results. Significant branches were defined after multiple-test correction using corrected p ≤ 0.05. In total, aBSREL detected 10 significant branches among the 141 tested, including lineage-specific signals in *T. cacao* Matina and *T. grandiflorum* C1074. The Malvaceae-putative lineage-specific clade did not show significant evidence of branch-level episodic diversifying selection after correction, supporting a predominantly conserved evolutionary pattern with localized episodes of diversification.

At the site level, MEME detected a limited number of codon positions with evidence of episodic diversifying selection (Supplementary Material 7). These sites were distributed across the alignment rather than concentrated in a single region. Some of these positions were located within the conserved COBRA domain, whereas others were located outside the domain. Because these sites were scattered and affected a limited subset of branches, they suggest localized sequence variation rather than broad adaptive divergence across the family.

Within the putative lineage-specific COBL clade, no branches showed significant evidence of episodic diversifying selection after multiple-test correction, including the representative genes TgrandC1074G00000008413.1 and TgrandC1074G00000008646.1. Together, these results indicate that, despite its phylogenetic distinctiveness, this group remains largely conserved and does not show evidence of widespread branch-level adaptive acceleration under the tests applied here.

## 4. Discussion

### 4.1. Conservation and diversification of *COBL* genes in *Theobroma* and *Herrania*

The comparative analysis of *COBL* genes in *T. grandiflorum*, *T. cacao*, and *H. umbratica* revealed a conserved gene family with moderate variation in copy number, genomic distribution, and structural organization. The identification of 62 *COBL* genes across the analyzed genomes is consistent with the broad conservation of this family in angiosperms, where COBL proteins have been associated with cell-wall organization, cellulose deposition, and developmental processes [18,20,23,64]. The presence of the COBRA domain in all retained proteins further supports their classification as COBL family members and indicates that the core domain architecture of this family is maintained in *Theobroma* and *Herrania*.

Although the COBL family was conserved, the gene number varied among genomes, with *T. cacao* Matina showing the largest repertoire. This variation appears to reflect the combined contribution of tandem, segmental, dispersed, and proximal duplications. Tandem duplications were frequent in *T. grandiflorum* and *T. cacao* Criollo, whereas segmental duplications were more represented in *T. cacao* Matina. Such patterns are consistent with the general role of gene duplication in shaping plant gene families and generating copy number variation [50;65-67]. These duplication patterns indicate that multiple genomic mechanisms contributed to *COBL* copy number variation in the analyzed genomes. Whether duplicated copies have acquired divergent regulatory or functional roles remains an open question, but the observed distribution provides a useful basis for prioritizing specific COBL groups in future comparative and functional studies.

Structural organization of *COBL* genes also supports the conservation of the two major subgroups previously described in plants [20]. Subgroup I genes generally displayed a more complex exon–intron organization, whereas Subgroup II genes were more compact, frequently containing two exons. This pattern agrees with previous studies in *A. thaliana*, maize, rice, cotton, and other plant species [18,20,23,27,64,68,69]. Therefore, despite copy number differences among the analyzed genomes, the main structural features of the COBL family appear to be conserved in *Theobroma* and *Herrania*.

### 4.2. A putative lineage-specific COBL group

A notable result of this study was the recovery of a putative lineage-specific COBL group containing representatives from the analyzed *Theobroma* and *Herrania* genomes. These sequences did not cluster directly with the *A. thaliana* COBL reference groups and were recovered within a Malvaceae/Byttnerioideae-associated clade in the expanded phylogenetic analysis. This pattern suggests that this group may represent a retained or diversified *COBL* lineage within the sampled Malvaceae. The recovery of this group in both *Theobroma* and *Herrania* suggests that it was likely present before the divergence of these analyzed lineages or was retained in parallel across closely related genomes.

Structural modeling showed that representative proteins from this putative lineage-specific group are broadly similar to COBL6-associated proteins, while sequence alignments revealed localized amino acid differences within the COBRA domain. These differences identify candidate residues for future investigation. The selection analyses reinforce this interpretation: the COBL family showed a predominant pattern of purifying selection, and the putative lineage-specific group did not show significant branch-level episodic diversifying selection after multiple-test correction. Thus, this group is best interpreted as a phylogenetically distinct but structurally conserved COBL lineage.

### 4.3. Functional implications and future directions

The biological relevance of *COBL* genes in plant cell-wall biology [20] makes this family a useful target for future studies in cupuassu, cacao, and related Malvaceae. *COBL4*-related genes are particularly interesting because this group was more represented in some genomes and has been associated with secondary cell wall organization, cellulose crystallinity, and mechanical strength in other species [23,69,70]. Similarly, *COBL7*, *COBL8*, and *COBL9*-related genes may be relevant candidates for studies on root and epidermal development, given their reported roles in root hair growth and cellulose deposition in *A. thaliana* [22]. However, these functional associations are based on orthology and literature from other species; they remain to be tested in *Theobroma* and *Herrania*.

## 5. Conclusion

Overall, this study provides a comparative framework for understanding the evolution and structural conservation of *COBL* genes in *Theobroma* and *Herrania*. The results support a model in which the COBL family is broadly conserved, shaped by multiple duplication processes, and characterized by subgroup-specific structural organization. The identification of a putative lineage-specific COBL group adds an additional layer of evolutionary interest, but its biological significance remains unresolved. Future work integrating tissue-specific expression, developmental transcriptomics, infection-related expression profiles, coexpression analyses, and functional validation will be necessary to determine whether specific *COBL* genes contribute to cell-wall-related traits such as tissue structure, fruit firmness, postharvest behavior, or pathogen responses in economically important Malvaceae crops.

## Supplementary Information

**Supplementary Material 1** All supplementary figures

**Supplementary Material 2** GenBank SRA accession numbers used for transcriptome assembly. (A) All *Theobroma cacao* RNA-Seq data used. (B) *Herrania umbratica* RNA-Seq used data. Source: Ray and Satya (2014).

**Supplementary Material 3** Plant species included in the expanded COBRA-like phylogenetic analysis. The table provides the taxonomic group, genome assembly accession, and number of predicted protein sequences containing the COBRA domain (PF04833.18) identified in each genome.

**Supplementary Material 4** Number of subgroups I and II COBRA-like family genes identified in *Theobroma cacao* (Criollo and Matina), *T. grandiflorum* (174 and 1074) and *Herrania umbratica*. The table includes information about subfamily, genes IDs, total size of proteins (in residues of amino acids, aa) and the region corresponding to the alignment with the COBRA domain (PFAM), indicating the starting and ending positions within the protein sequence.

**Supplementary Material 5** TPM (Transcripts Per Million) values for *COBRA-like* genes in *T. cacao* (Criollo and Matina), *T. grandiflorum* (174 and 1074), and *Herrania umbratica*.

**Supplementary Material 6** Assessment of the structural quality of proteins from the COBL6, COBL7,8,9, COBL10,11 groups from *Theobroma* and *Herrania*. The table presents the ERRAT (Overall Quality Factor) values, which indicate the general quality of the structural model, and the PROCHECK results (Ramachandran plot, %), which show the distribution of residues in the conformationally permitted regions: core (favored), allow, gener (marginal) and disall (disallowed).

**Supplementary Material 7** Site-level episodic diversifying selection inferred by MEME for the COBRA-Like gene family. The table reports codon positions from the codon-aware alignment used in the Hyphy/MEME analysis, together with synonymous substitution rates (α), non-synonymous rate classes under the MEME model (β-and β+), their corresponding weights (p- and p+), likelihood ratio test values (LRT), p- values, number of branches inferred to be under episodic selection, total branch length, and log-likelihood estimates from MEME and FEL. Codon positions refer to the multiple sequence alignment and therefore do not necessarily correspond to absolute amino acid positions in each individual protein. Sites with p ≤ 0.05 were considered significant for episodic diversifying selection. FEL estimates are provided as complementary site-level substitution parameters.

## Funding and acknowledgments

This study was financed in part by the Coordenação de Aperfeiçoamento de Pessoal de Nível Superior – Brasil (CAPES), Finance Code 001. We thank CAPES for the doctoral scholarship and acknowledge the support provided by UNESP and the Graduate Program in Agronomy (Genetics and Plant Breeding) (process number 88887.615107/2021-00). We also thank Embrapa Amazônia Oriental for providing the samples. This study was also funded by the Fundação de Amparo à Pesquisa do Estado de São Paulo (FAPESP), grants #2019/25176-0 to AMV, #2024/05163-0 to JVAA, #2023/10314-4 to VT, #2023/08702-6 to ASB, and #2023/04372-1 to MMO. AMV and VT are affiliated with the Center for Research on Biodiversity Dynamics and Climate Change (CEPID-FAPESP, grant #2021/10639–5).

## Author contributions

**Rafaely Pantoja Oliveira:** Writing – review & editing, Writing – original draft, Validation, Methodology, Formal analysis, Data curation, Conceptualization. **João Victor dos Anjos Almeida:** Writing – review & editing, Methodology, Formal analysis, Data curation. **Vitor Trinca:** Writing – review & editing, Methodology, Formal analysis, Data curation. **Anelise Stella Ballaben:** Writing – review & editing, Data curation. **Mauro de Medeiros Oliveira:** Writing – review & editing, Validation, Methodology, Formal analysis, Data curation. **Alessandro de Mello Varani:** Writing – review & editing, Supervision, Resources, Project administration, Funding acquisition, Conceptualization.

## Declarations

### Ethics approval and consent to participate

Not applicable.

### Consent for publication

Not applicable.

### Data availability

Genomic sequences and annotation files analyzed in this study were retrieved from public databases under the accession numbers PRJNA383741, PRJEB14326, and PRJNA239715. Additional publicly available genomic datasets used in comparative analyses are described throughout the manuscript and supplementary material.

## REFERENCES

1. Colli-Silva M, Richardson JE, Pirani JR. A taxonomic dataset of preserved specimen occurrences of Theobroma and Herrania (Malvaceae, Byttnerioideae) stored in 2020. Biodivers Data J. 2023;11:e99646. 10.3897/BDJ.11.e99646.

2. Colli-Silva M, Richardson JE, Neves EG, Watling J, Figueira A, Pirani JR. Domestication of the Amazonian fruit tree cupuaçu may have stretched over the past 8000 years. Commun Earth Environ. 2023;4(1):401. 10.1038/s43247-023-01066-z.

3. Flora e Funga do Brasil, 2024. Jardim Botânico do Rio de Janeiro. 2024. Available from: https://floradobrasil.jbrj.gov.br/consulta/#CondicaoTaxonCP. Accessed May 04, 2026.

4. Bletter N, Daly DC. Cacao and its relatives in South America: An overview of taxonomy, ecology, biogeography, chemistry, and ethnobotany. In: Chocolate in Mesoamerica: A Cultural History of Cacao. 2006. p. 31–68. 10.5744/florida/9780813029535.003.0002.17.

5. Sousa Silva CR, Figueira A. Phylogenetic analysis of Theobroma (Sterculiaceae) based on Kunitz-like trypsin inhibitor sequences. Plant Syst Evol. 2005;250(1):93– 104. 10.1007/s00606-004-0223-2.

6. Cohen KDO, Jackix MDNH. Estudo do liquor de cupuaçu. Food Sci Technol. 2005;25(1):182–90. 10.1590/S0101-20612005000100030.

7. Alves RM, Sebbenn AM, Artero AS, Clement C, Figueira A. High levels of genetic divergence and inbreeding in populations of cupuassu (Theobroma grandiflorum). Tree Genet Genomes. 2007;3(4):289–98. 10.1007/s11295-006-0066-9.

8. Genovese MI, Lannes SCDS. Comparison of total phenolic content and antiradical capacity of powders and “chocolates” from cocoa and cupuassu. Food Sci Technol. 2009;29:810–4. 10.1590/S0101-20612009000400017.

9. McNeil M. Chocolate in Mesoamerica: A cultural history of cacao. University Press of Florida. 2006

10. Silva da Costa R, Pinheiro WBDS, Arruda MSP, Costa CEF, Converti A, Ribeiro Costa RM, Silva Júnior JOC. Thermoanalytical and phytochemical study of the cupuassu (Theobroma grandiflorum Schum.) seed by-product in different processing stages. J Therm Anal Calorim. 2022;147(1):275–84. 10.1007/s10973-020-10347-0.

11. Benchimol RL, Silva CM, Melo BVA. O cupuaçuzeiro na Amazônia: ciência, tecnologia e produção. Embrapa. Available from: https://www.alice.cnptia.embrapa.br/alice/bitstream/doc/1175355/1/LV-CupuacueiroAmazonia.pdf. Accessed May 4, 2026.

12. Secretaria de Estado de Desenvolvimento Agropecuário e da Pesca (SEDAP). SEDAP – Secretaria de Estado de Desenvolvimento Agropecuário e da Pesca. 2024. Available from: https://sedap.pa.gov.br/boletim-cvis. Accessed May 04, 2026.

13. Alves RM, Chaves SFDS. Selection of Theobroma grandiflorum clones adapted to agroforestry systems using an additive index. Acta Sci Agron. 2023;45:e57519. 10.4025/actasciagron.v45i1.57519.

14. Alves RM, Silva CRDS, Albuquerque PSBD, Santos VSD. Phenotypic and genotypic characterization and compatibility among genotypes to select elite clones of cupuassu. Acta Amaz. 2017;47:175–84. 10.1590/1809-4392201602104.

15. De Abreu VA, Alves RM, Silva SR, Ferro JA, Domingues DS, Miranda VF, Varani AM. Comparative analyses of Theobroma cacao and T. grandiflorum mitogenomes reveal conserved gene content embedded within complex and plastic structures. Gene. 2023;849:146904. 10.1016/j.gene.2022.146904.

16. Mournet P, de Albuquerque PSB, Alves RM, Silva-Werneck JO, Rivallan R, Marcellino LH, Clément D. A reference high-density genetic map of Theobroma grandiflorum (Willd. ex Spreng) and QTL detection for resistance to witches’ broom disease (Moniliophthora perniciosa). Tree Genet Genomes. 2020;16(6):89. 10.1007/s11295-020-01479-3.

17. Niu YF, Ni SB, Liu J. The complete chloroplast genome of Theobroma grandiflorum, an important tropical crop. Mitochondrial DNA B Resour. 2019;4(2):4157–8. 10.1080/23802359.2019.1693291.18.

18. Li S, Ge FR, Xu M, Zhao XY, Huang GQ, Zhou LZ, Wang JG, Kombrink A, McCormick S, Zhang XS, Zhang Y. Arabidopsis COBRA-LIKE 10, a GPI-anchored protein, mediates directional growth of pollen tubes. Plant J. 2013;74(3):486–97. 10.1111/tpj.12139.

19. Yang Q, Wang S, Chen H, You L, Liu F, Liu Z. Genome-wide identification and expression profiling of the COBRA-like genes reveal likely roles in stem strength in rapeseed (Brassica napus L.). PLOS ONE. 2021;16(11):e0260268. 10.1371/journal.pone.0260268.

20. Roudier F, Schindelman G, DeSalle R, Benfey PN. The COBRA family of putative GPI-anchored proteins in Arabidopsis. A new fellowship in expansion. Plant Physiol. 2002;130(2):538–48. 10.1104/pp.007468.

21. Ko JH, Kim JH, Jayanty SS, Howe GA, Han KH. Loss of function of COBRA, a determinant of oriented cell expansion, invokes cellular defence responses in Arabidopsis thaliana. J Exp Bot. 2006;57(12):2923–36. 10.1093/jxb/erl052.

22. Li Z, Zhou T, Sun P, Chen X, Gong L, Sun P, Ge S, Liang YK. COBL9 and COBL7 synergistically regulate root hair tip growth via controlling apical cellulose deposition. Biochem Biophys Res Commun. 2022;596:6–13. 10.1016/j.bbrc.2022.01.096.

23. Brady SM, Song S, Dhugga KS, Rafalski JA, Benfey PN. Combining expression and comparative evolutionary analysis. The COBRA gene family. Plant Physiol. 2007;143(1):172–87. 10.1104/pp.106.087262.

24. Li Y, Qian Q, Zhou Y, Yan M, Sun L, Zhang M, Fu Z, Wang Y, Han B, Pang X, Chen M, Li J. BRITTLE CULM1, which encodes a COBRA-like protein, affects the mechanical properties of rice plants. Plant Cell. 2003;15(9):2020–31. 10.1105/tpc.011775.

25. Dai X, You C, Wang L, Chen G, Zhang Q, Wu C. Molecular characterization, expression pattern, and function analysis of the OsBC1L family in rice. Plant Mol Biol. 2009;71(4):469–81. 10.1007/s11103-009-9537-3.

26. Cao Y, Tang X, Giovannoni J, Xiao F, Liu Y. Functional characterization of a tomato COBRA-like gene functioning in fruit development and ripening. BMC Plant Biol. 2012;12(1):211. 10.1186/1471-2229-12-211.

27. Niu E, Shang X, Cheng C, Bao J, Zeng Y, Cai C, Du X, Guo W.Comprehensive analysis of the COBRA-like (COBL) gene family in Gossypium identifies two COBLs potentially associated with fiber quality. PLOS ONE. 2015;10(12):e0145725. 10.1371/journal.pone.0145725.

28. Ye X, Kang BG, Osburn LD, Cheng ZM. The COBRA gene family in Populus and gene expression in vegetative organs and in response to hormones and environmental stresses. Plant Growth Regul. 2009;58(2):211–23. 10.1007/s10725-009-9369-9.

29. Alves RM, Abreu VAC, Oliveira RP, Almeida JVA, Oliveira MM, Silva SR, Paschoal AR, Almeida SS, Souza PAF, Ferro JA, Miranda VFO, Figueira A, Domingues DS, Varani AM. Genomic decoding of Theobroma grandiflorum (cupuassu) at chromosomal scale: evolutionary insights for horticultural innovation. GigaScience. 2024;13:giae027. 10.1093/gigascience/giae027.

30. Abreu VAC, Alves RM, Oliveira MM, Trinca V, Falcão LL, Marcellino LH, Figueira A, Domingues DS, Varani A. Integrative chromosome-scale genome analysis of cupuassu provides insights into witches’ broom disease resistance and expands genomic resources for Theobroma. Plant Genome. 2026;19(1):e70196. 10.1002/tpg2.70196.

31. Alves RM, Resende MDVD. Genetic evaluation of individuals and progenies of Theobroma grandiflorum in the state of Pará and estimates of genetic parameters. Rev Bras Frutic. 2008;30:696–701. 10.1590/S0100-29452008000300023.

32. Argout X, Salse J, Aury JM, Guiltinan MJ, Droc G, Gouzy J, Allegre M, Chaparro C, Legavre T, Maximova SN, Abrouk M, Murat F, Fouet O, Poulain J, Ruiz M, Roguet Y, Rodier-Goud M, Barbosa-Neto JF, Sabot F, Kudrna D, Ammiraju JSS, Schuster SC, Carlson JE, Sallet E, Schiex T, Dievart A, Kramer M, Gelley L, Shi Z, Bérard A, Viot C, Boccara M, Risterucci AM, Guignon V, Sabau X, Axtell MJ, Ma Z, Zhang Y, Brown S, Bourge M, Golser W, Song X, Clement D, Rivallan R, Tahi M, Akaza JM, Pitollat B, Gramacho K, D’Hont A, Brunel D, Infante D, Kebe I, Costet P, Wing R, McCombie WR, Guiderdoni E, Quetier F, Panaud O, Wincker P, Bocs S, Lanaud C. The genome of Theobroma cacao. Nat Genet. 2011;43(2):101–8. 10.1038/ng.736.

33. Motamayor JC, Mockaitis K, Schmutz J, Haiminen N, Livingstone D III, Cornejo O, Findley SD, Zheng P, Utro F, Royaert S, Saski C, Jenkins J, Podicheti R, Zhao M, Scheffler BE, Stack JC, Feltus FA, Mustiga GM, Amores F, Phillips W, Marelli JP, May GD, Shapiro H, Ma J, Bustamante CD, Schnell RJ, Main D, Gilbert D, Parida L, Kuhn DN. The genome sequence of the most widely cultivated cacao type and its use to identify candidate genes regulating pod color. Genome Biol. 2013;14(6):1–25. 10.1186/gb-2013-14-6-r53.

34. Roudier F, Fernandez AG, Fujita M, Himmelspach R, Börner GHH, Schindelman G, Song S, Baskin TI, Dupree P, Wasteneys GO, Benfey PN. COBRA, an Arabidopsis extracellular glycosyl-phosphatidyl inositol-anchored protein, specifically controls highly anisotropic expansion through its involvement in cellulose microfibril orientation. Plant Cell. 2005;17(6):1749–63. 10.1105/tpc.105.031732.

35. Hochholdinger F, Wen TJ, Zimmermann R, Chimot-Marolle P, Costa e Silva O, Bruce W, Lamkey KR, Wienand U, Schnable PS. The maize (Zea mays L.) roothairless3 gene encodes a putative GPI-anchored, monocot-specific, COBRA-like protein that significantly affects grain yield. Plant J. 2008;54(5):888–98. 10.1111/j.1365-313x.2008.03459.x.

36. Ben-Tov D, Abraham Y, Stav S, Thompson K, Loraine A, Elbaum R, Souza A, Pauly M, Kieber JJ, Harpaz-Saad S. COBRA-LIKE2, a member of the glycosylphosphatidylinositol-anchored COBRA-LIKE family, plays a role in cellulose deposition in Arabidopsis seed coat mucilage secretory cells. Plant Physiol. 2015;167(3):711–24. 10.1104/pp.114.240671.

37. Ben-Tov D, Idan-Molakandov A, Hugger A, Ben-Shlush I, Günl M, Yang B, Usadel B, Harpaz-Saad S. The role of COBRA-LIKE 2 function, as part of the complex network of interacting pathways regulating Arabidopsis seed mucilage polysaccharide matrix organization. Plant J. 2018;94(3):497–512. 10.1111/tpj.13871.

38. Niu E, Fang S, Shang X, Guo W. Ectopic expression of GhCOBL9A, a cotton glycosyl-phosphatidyl inositol-anchored protein encoding gene, promotes cell elongation, thickening and increased plant biomass in transgenic Arabidopsis. Mol Genet Genomics. 2018;293(5):1191–204. 10.1007/s00438-018-1452-3.

39. Brown DM, Zeef LA, Ellis J, Goodacre R, Turner SR. Identification of novel genes in Arabidopsis involved in secondary cell wall formation using expression profiling and reverse genetics. Plant Cell. 2005;17(8):2281–95. 10.1105/tpc.105.031542.

40. Ko JH, Kim JH, Jayanty SS, Howe GA, Han KH. Loss of function of COBRA, a determinant of oriented cell expansion, invokes cellular defence responses in Arabidopsis thaliana. J Exp Bot. 2006;57(12):2923–36. 10.1111/tpj.12139.

41. Hok S, Danchin EGG, Allasia V, Panabières F, Attard A, Keller H. An Arabidopsis (malectin-like) leucine-rich repeat receptor-like kinase contributes to downy mildew disease. Plant Cell Environ. 2011;34(11):1944–57. 10.1111/j.1365-3040.2011.02390.x.

42. Schaaf G, Honsbein A, Meda AR, Kirchner S, Wipf D, von Wirén N. AtIREG2 encodes a tonoplast transport protein involved in iron-dependent nickel detoxification in Arabidopsis thaliana roots. J Biol Chem. 2006;281(35):25532–40. 10.1074/jbc.m601062200.

43. Ge S, Sun P, Wu W, Chen X, Wang Y, Zhang M, Huang J, Liang YK. COBL7 is required for stomatal formation via regulation of cellulose deposition in Arabidopsis. New Phytol. 2024;241(1):227–42. 10.1111/nph.19327.

44. Jones MA, Raymond MJ, Smirnoff N. Analysis of the root-hair morphogenesis transcriptome reveals the molecular identity of six genes with roles in root-hair development in Arabidopsis. Plant J. 2006;45(1):83–100. 10.1111/j.1365-313x.2005.02609.x.

45. Li H, Yang Y, Zhang H, Li C, Du P, Bi M, Chen T, Qian D, Niu Y, Ren H, An L, Xiang Y. The Arabidopsis GPI-anchored protein COBL11 is necessary for regulating pollen tube integrity. Cell Rep. 2023;42(11):113353. 10.1016/j.celrep.2023.113353.

46. Potter SC, Luciani A, Eddy SR, Park Y, Lopez R, Finn RD. HMMER webserver: 2018 update. Nucleic Acids Res. 2018;46(W1):W200–4. 10.1093/nar/gky448.

47. Mistry J, Chuguransky S, Williams L, Qureshi M, Salazar GA, Sonnhammer ELL, Tosatto SCE, Paladin L, Raj S, Richardson LJ, Finn RD, Bateman A. Pfam: The protein families database in 2021. Nucleic Acids Res. 2021;49(D1):D412–9. 10.1093/nar/gkaa913.

48. Chao J, Li Z, Sun Y, Aluko OO, Wu X, Wang Q, Liu G. MG2C: A user-friendly online tool for drawing genetic maps. Mol Hortic. 2021;1(1):1–4. 10.1186/s43897-021-00020-x.

49. Doyle JJ, Doyle JL. A rapid DNA isolation procedure for small quantities of fresh leaf tissue. Phytochem Bull. 1987;19:11–5. https://webpages.charlotte.edu/∼jweller2/pages/BINF8350f2011/BINF8350_Readings/Doyle_plantDNAextractCTAB_1987.pdf

50. Wang Y, Tang H, DeBarry JD, Tan X, Li J, Wang X, Lee TH, Jin H, Marler B, Guo H, Kissinger JC, Paterson AH. MCScanX: a toolkit for detection and evolutionary analysis of gene synteny and collinearity. Nucleic Acids Res. 2012;40(7):e49. 10.1093/nar/gkr1293.

51. Chen C, Wu Y, Li J, Wang X, Zeng Z, Xu J, Liu Y, Feng J, Chen H, He Y, Xia R. TBtools-II: A “one for all, all for one” bioinformatics platform for biological big-data mining. Mol Plant. 2023;16(11):1733–42. 10.1016/j.molp.2023.09.010.

52. Trifinopoulos J, Nguyen LT, von Haeseler A, Minh BQ. W-IQ-TREE: a fast online phylogenetic tool for maximum likelihood analysis. Nucleic Acids Res. 2016;44(W1):W232–5. 10.1093/nar/gkw256.

53. Katoh K, Standley DM. MAFFT multiple sequence alignment software version 7: improvements in performance and usability. Mol Biol Evol. 2013;30:772–80. 10.1093/molbev/mst010.

54. Nguyen LT, Schmidt HA, von Haeseler A, Minh BQ. IQ-TREE: a fast and effective stochastic algorithm for estimating maximum-likelihood phylogenies. Mol Biol Evol. 2015;32:268–74. 10.1093/molbev/msu300.

55. Kalyaanamoorthy S, Minh BQ, Wong TK, von Haeseler A, Jermiin LS. ModelFinder: fast model selection for accurate phylogenetic estimates. Nat Methods. 2017;14:587– 9. 10.1038/nmeth.4285.

56. Letunic I, Bork P. Interactive Tree of Life (iTOL) v6: recent updates to the phylogenetic tree display and annotation tool. Nucleic Acids Res. 2024;52(W1):W78– 82. 10.1093/nar/gkae268.

57. Quevillon E, Silventoinen V, Pillai S, Harte N, Mulder N, Apweiler R, Lopez R. InterProScan: protein domains identifier. Nucleic Acids Res. 2005;33(suppl_2):W116–20. 10.1093/nar/gki442.

58. Suyama M, Torrents D, Bork P. PAL2NAL: robust conversion of protein sequence alignments into the corresponding codon alignments. Nucleic Acids Res. 2006;34(suppl_2):W609–12. 10.1093/nar/gkl315.

59. Smith MD, Wertheim JO, Weaver S, Murrell B, Scheffler K, Kosakovsky Pond SL. Less is more: an adaptive branch-site random effects model for efficient detection of episodic diversifying selection. Mol Biol Evol. 2015;32:1342–53. 10.1093/molbev/msv022.

60. Murrell B, Wertheim JO, Moola S, Weighill T, Scheffler K, Kosakovsky Pond SL. Detecting individual sites subject to episodic diversifying selection. PLOS Genetics. 2012;8(7):e1002764. 10.1371/journal.pgen.1002764.

61. Abramson J, Adler J, Dunger J, Evans R, Green T, Pritzel A, Ronneberger O, Willmore L, Ballard AJ, Bambrick J, Bodenstein SW, Evans DA, Hung CC, O’Neill M, Reiman D, Tunyasuvunakool K, Wu Z, Žemgulytė A, Arvaniti E, Beattie C, Bertolli O, Bridgland A, Cherepanov A, Congreve M, Cowen-Rivers AI, Cowie A, Figurnov M, Fuchs FB, Gladman H, Jain R, Khan YA, Low CMR, Perlin K, Potapenko A, Savy P, Singh S, Stecula A, Thillaisundaram A, Tong C, Yakneen S, Zhong ED, Zielinski M, Žídek A, Bapst V, Kohli P, Jaderberg M, Hassabis D, Jumper JM. Accurate structure prediction of biomolecular interactions with AlphaFold 3. Nature. 2024;630:493–500. 10.1038/s41586-024-07487-w.

62. Ramachandran GN, Ramakrishnan C, Sasisekharan V. Stereochemistry of polypeptide chain configurations. J Mol Biol. 1963;7:95–99. 10.1016/S0022-2836(63)80023-6.

63. Li Z, Jaroszewski L, Iyer M, Sedova M, Godzik A. FATCAT 2.0: towards a better understanding of the structural diversity of proteins. Nucleic Acids Res. 2020;48:W60–4. 10.1093/nar/gkaa443.

64. Sangi S, Araújo PM, Coelho FS, Gazara RK, Almeida-Silva F, Venancio TM, Grativol C. Genome-wide analysis of the COBRA-like gene family supports gene expansion through whole-genome duplication in soybean (Glycine max). Plants. 2021;10:167. 10.3390/plants10010167.

65. Hanada K, Zou C, Lehti-Shiu MD, Shinozaki K, Shiu SH. Importance of lineage-specific expansion of plant tandem duplicates in the adaptive response to environmental stimuli. Plant Physiol. 2008;148:993–1003. 10.1104/pp.108.122457.

66. Zou C, Lehti-Shiu MD, Thomashow M, Shiu SH. Evolution of stress-regulated gene expression in duplicate genes of Arabidopsis thaliana. PLOS Genetics. 2009;5(7):e1000581. 10.1371/journal.pgen.1000581.

67. Wang J, Tao F, Marowsky NC, Fan C. Evolutionary fates and dynamic functionalization of young duplicate genes in Arabidopsis genomes. Plant Physiol. 2016;172:427–40. 10.1104/pp.16.01177.

68. Ching A, Dhugga KS, Appenzeller L, Meeley R, Bourett TM, Howard RJ, Rafalski A. Brittle stalk 2 encodes a putative glycosylphosphatidylinositol-anchored protein that affects mechanical strength of maize tissues by altering the composition and structure of secondary cell walls. Planta. 2006;224:1174–84. 10.1007/s00425-006-0299-8.

69. Li P, Liu Y, Tan W, Chen J, Zhu M, Lv Y, Liu Y, Yu S, Zhang W, Cai H. Brittle Culm 1 encodes a COBRA-like protein involved in secondary cell wall cellulose biosynthesis in Sorghum. Plant Cell Physiol. 2019;60:788–801. 10.1093/pcp/pcy246.

70. Schindelman G, Morikami A, Jung J, Baskin TI, Carpita NC, Derbyshire P, McCann MC, Benfey PN. COBRA encodes a putative GPI-anchored protein, which is polarly localized and necessary for oriented cell expansion in Arabidopsis. Genes Dev. 2001;15:1115–27.. 10.1101/gad.879101.

