## Supplementary_Material_1 for "Comparative Genomics of COBRA-like Genes in *Theobroma* and *Herrania* Reveals Structural Conservation and Lineage-Specific Variation"

**Supplementary Fig. 1** Molecular modeling of COBL proteins from groups COBL6, COBL7,8,9, and COBL10,11, compared to the putative lineage-specific COBL group identified in this study in Amazonian Malvaceae species. The Ramachandran plot was generated using the PROCHECK program for the proteins. The plot presents the distribution of protein residues in different conformational regions based on the $\phi$ and $\psi$ angles: most favored regions (red), additionally allowed regions (yellow), generously allowed regions (light yellow), and disallowed regions (white). These regions reflect the stability and energetic feasibility of peptide conformations.

**Supplementary Fig. 2** Domain amino acid sequence alignment COBRA in proteins COBL6 (blue rectangle) and putative lineage-specific COBL group (orange rectangle). The analysis highlights specific changes in the amino acid groups that make up the functional domain, with changes between amino acids of distinct chemical properties indicated by red asterisks, revealing structural variations that may be associated with adaptive functions in the Amazonian Malvaceae.

**Supplementary Fig. 3** Amino acid sequence alignment of the COBRA domain (PFAM: PF04833) against the proteins COBL1, 2, 3, 4, 5, 6, 7, 8, 9, 10, 11, 12, and an putative lineage-specific COBL group. The analysis reveals specific changes in the amino acids that compose the functional domain, highlighting structural variations that may be associated with adaptive functions in Amazonian Malvaceae.

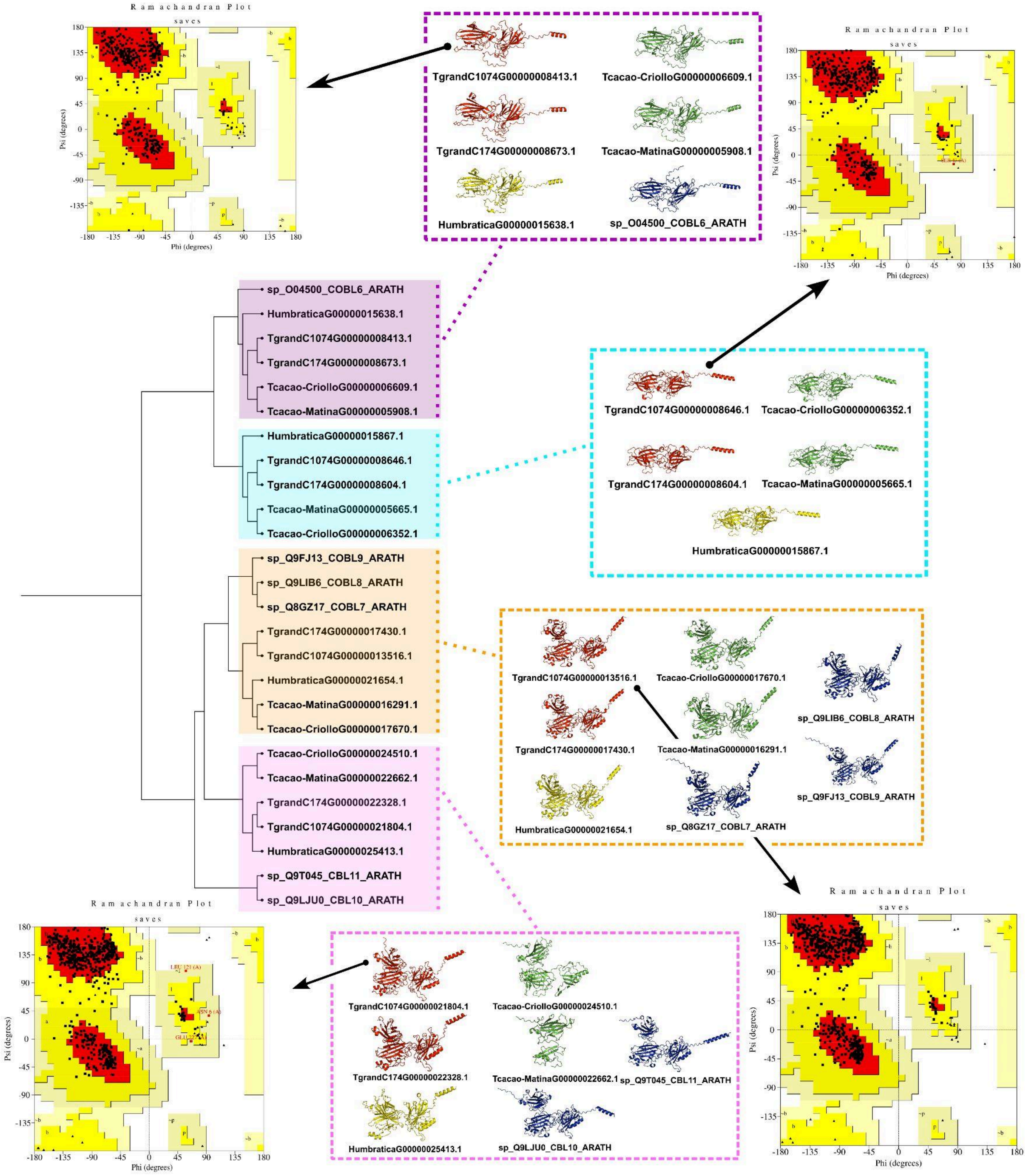


Supplementary Fig. 1 Molecular modeling of COBL proteins from groups COBL6, COBL7,8,9, and COBL10,11, compared to the putative lineage-specific COBL group identified in this study in Amazonian Malvaceae species. The Ramachandran plot was generated using the PROCHECK program for the proteins. The plot presents the distribution of protein residues in different conformational regions based on the ϕ and ψ angles: most favored regions (red), additionally allowed regions (yellow), generously allowed regions (light yellow), and disallowed regions (white). These regions reflect the stability and energetic feasibility of peptide conformations.

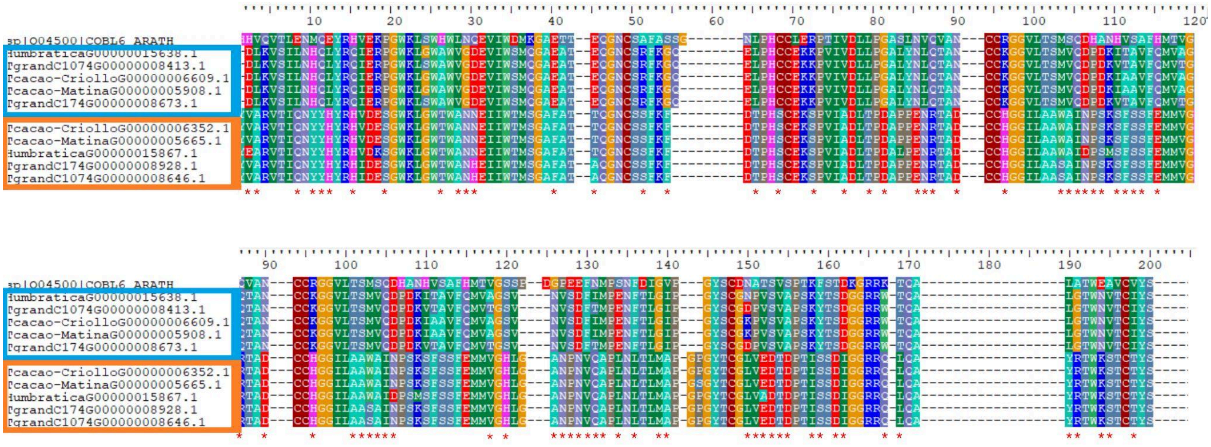

Supplementary Fig. 2 Domain amino acid sequence alignment COBRA in proteins COBL6 (blue rectangle) and putative lineage-specific COBL group (orange rectangle). The analysis highlights specific changes in the amino acid groups that make up the functional domain, with changes between amino acids of distinct chemical properties indicated by red asterisks, revealing structural variations that may be associated with adaptive functions in the Amazonian Malvaceae.

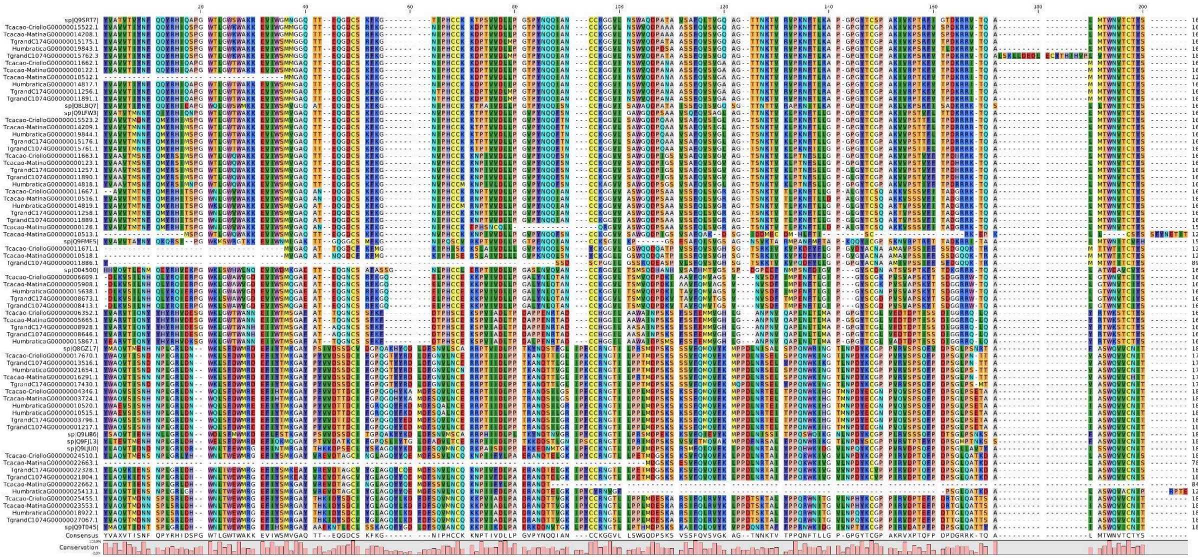

Supplementary Fig. 3 Amino acid sequence alignment of the COBRA domain (PFAM: PF04833) against the proteins COBL1, 2, 3, 4, 5, 6, 7, 8, 9, 10, 11, 12, and an putative lineage-specific COBL group. The analysis reveals specific changes in the amino acids that compose the functional domain, highlighting structural variations that may be associated with adaptive functions in Amazonian Malvaceae.
